## Supplementary Data for "H1 restricts euchromatin-associated methylation pathways from heterochromatic encroachment"

Supplementary Figures 1-8

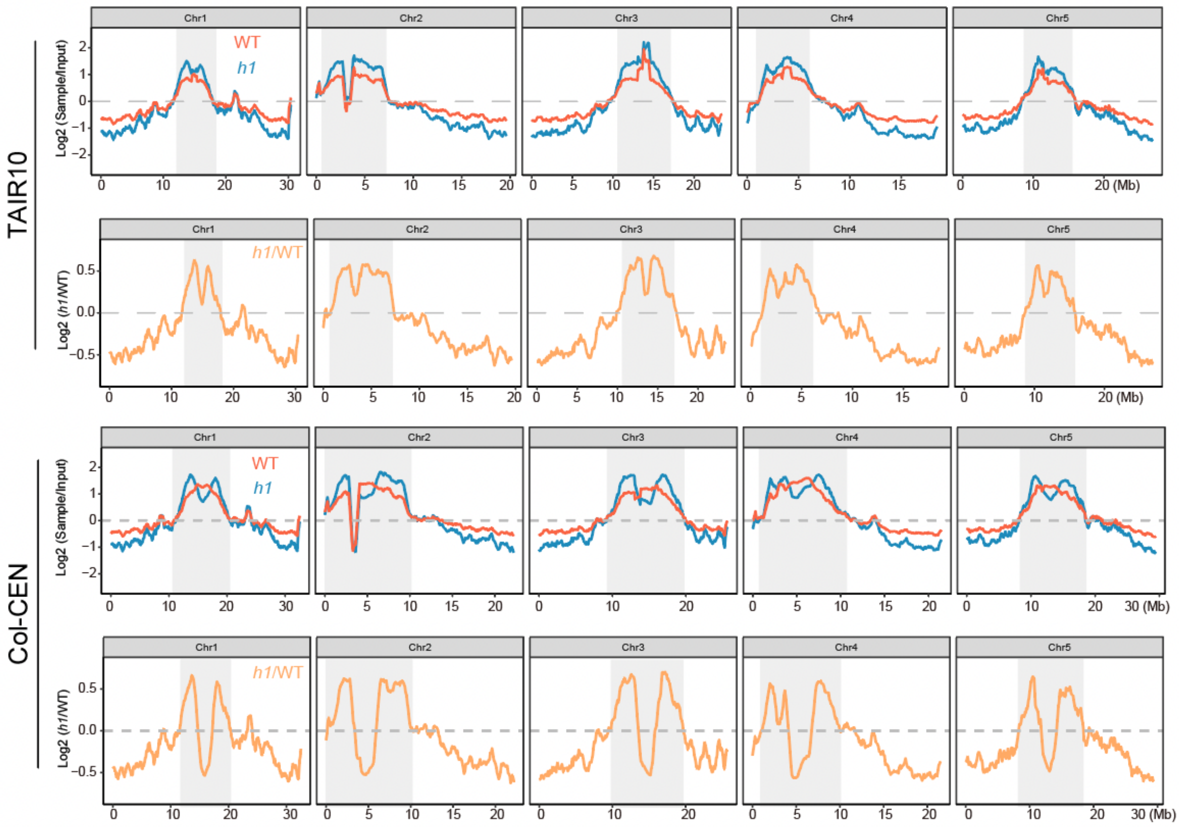


**Supplementary Figure 1:** NRPE1 enrichment in WT vs *h1* when mapped to either TAIR10 or Col-CEN genome assemblies (Naish et al., 2021).

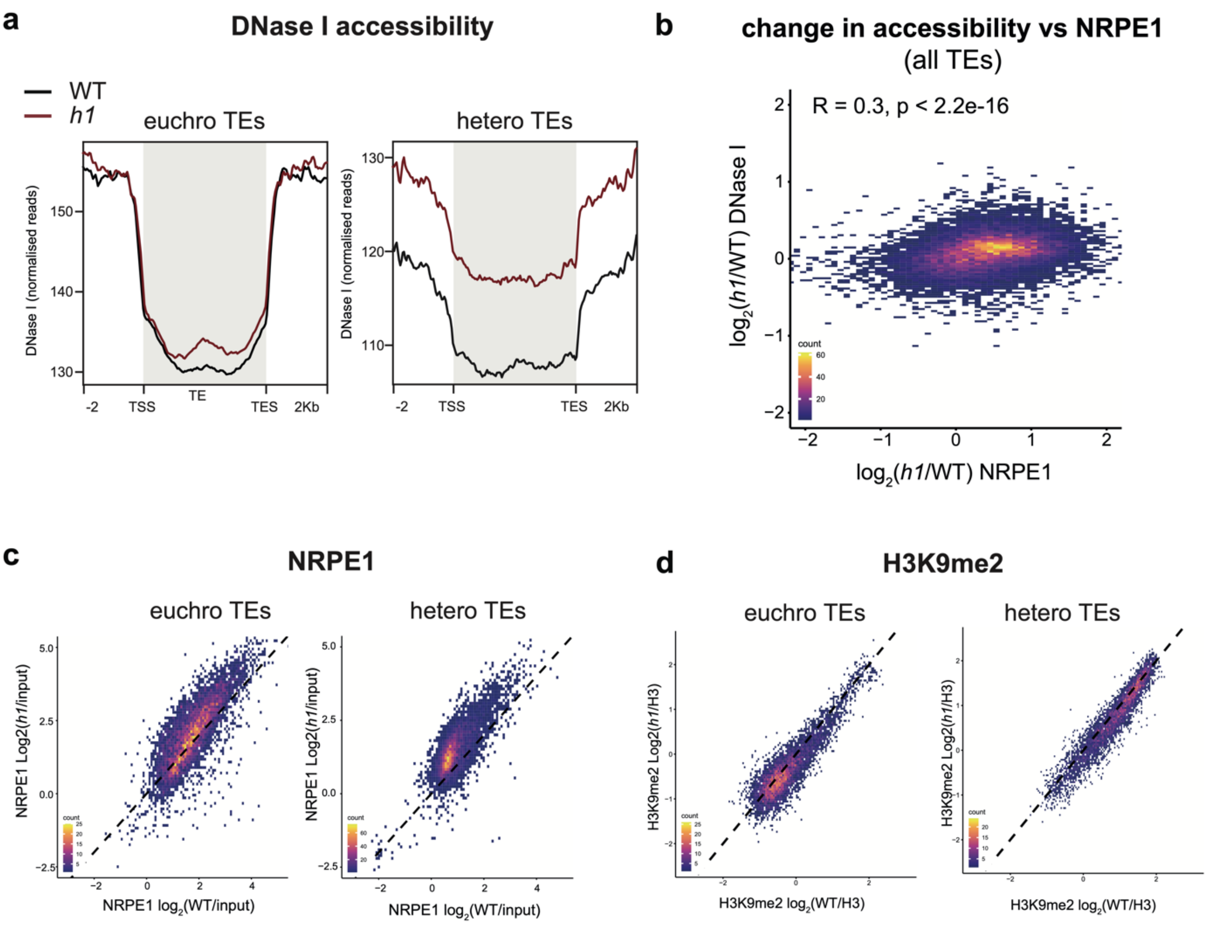


**Supplementary Figure 2:** A) Metaplot of DNase I accessibility in WT vs *h1* over euchromatic vs heterochromatic TEs. Data is obtained from (Choi et al., 2020). B) Scatterplot showing correlation between change in accessibility and NRPE1 occupancy in *h1*. R squared and p-value are indicated. C) Scatterplot correlation comparing NRPE1 enrichment in WT vs *h1* at euchromatic versus heterochromatic TEs. D) Scatterplot correlation comparing H3K9me2 enrichment in WT vs *h1* at euchromatic versus heterochromatic TEs.

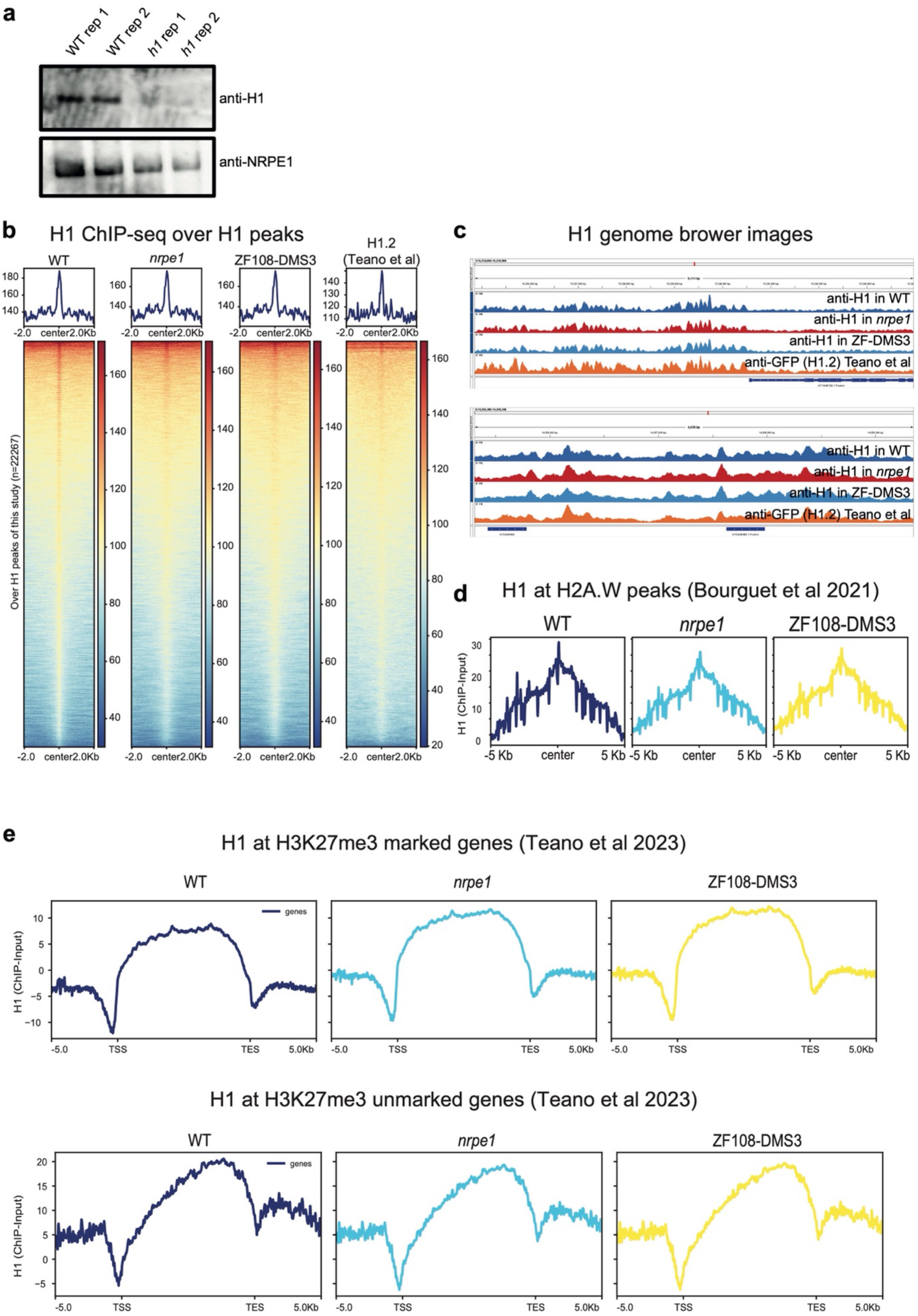


**Supplementary Figure 3:** A) Western blot of H1 and NRPE1, using same antibodies as used for ChIPseq. H1 is only detectable in the WT samples, not *h1* mutant. B) H1 ChIP-seq metaplots with over

WT H1 peaks with libraries indicated C) Genome browser images of H1 ChIP-seq track D) H1 enrichment over H2A.W peaks in euchromatic arms (as defined in Bourguet et al 2021). E) H1 enrichment patterns over H3K27me3 marked and unmarked genes (as defined in Teano et al 2023).

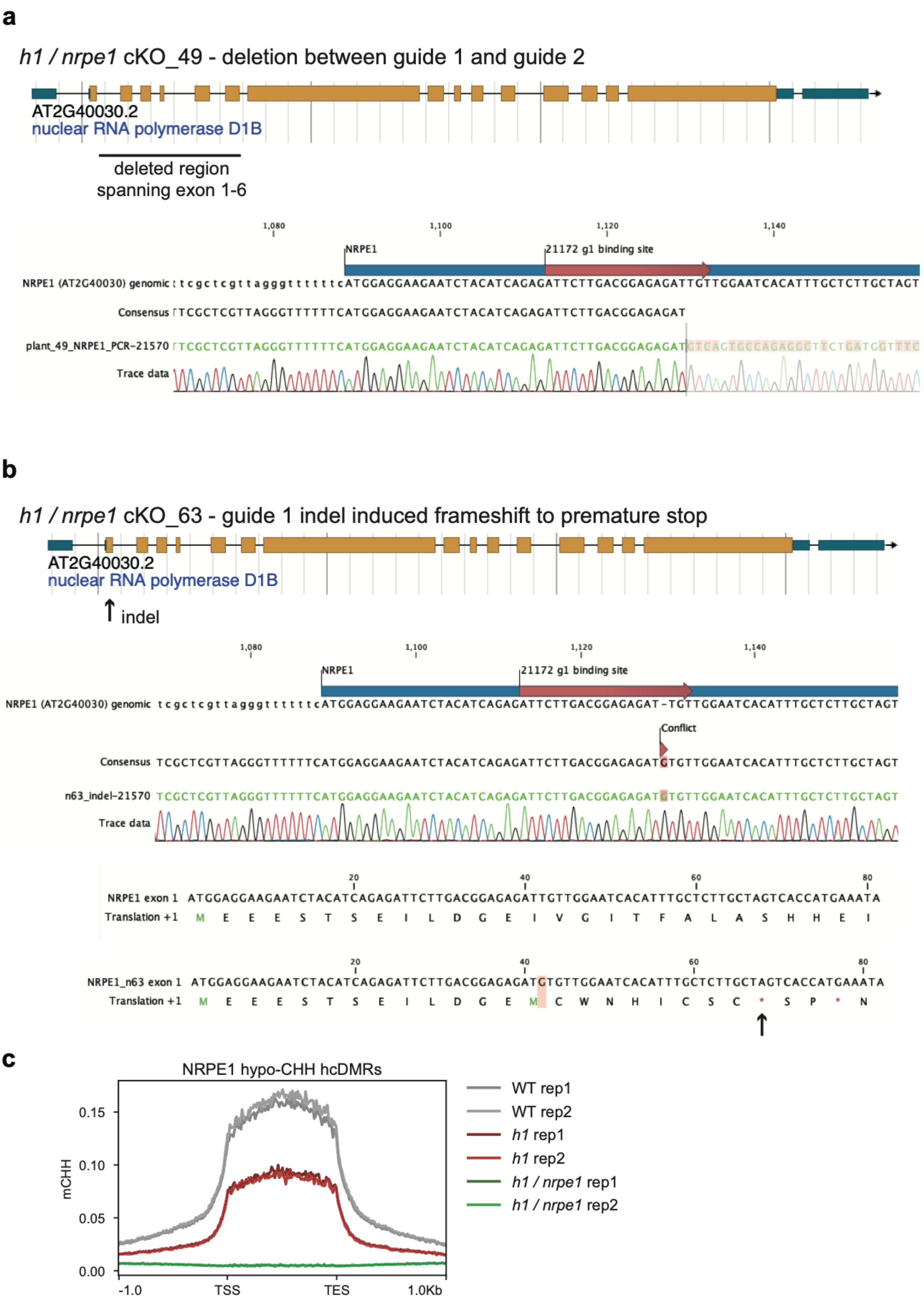


**Supplementary Figure 4**: NRPE1 CRISPR knock outs. A) *h1* / *nrpe1* cKO_49 has a deletion spanning exons 1-6. B) *h1* / *nrpe1* cKO_63 has an indel induced frameshift leading to a premature stop codon in exon 1. C) Both knockouts display complete loss of CHH methylation at NRPE1 hypo CHH DMRs.

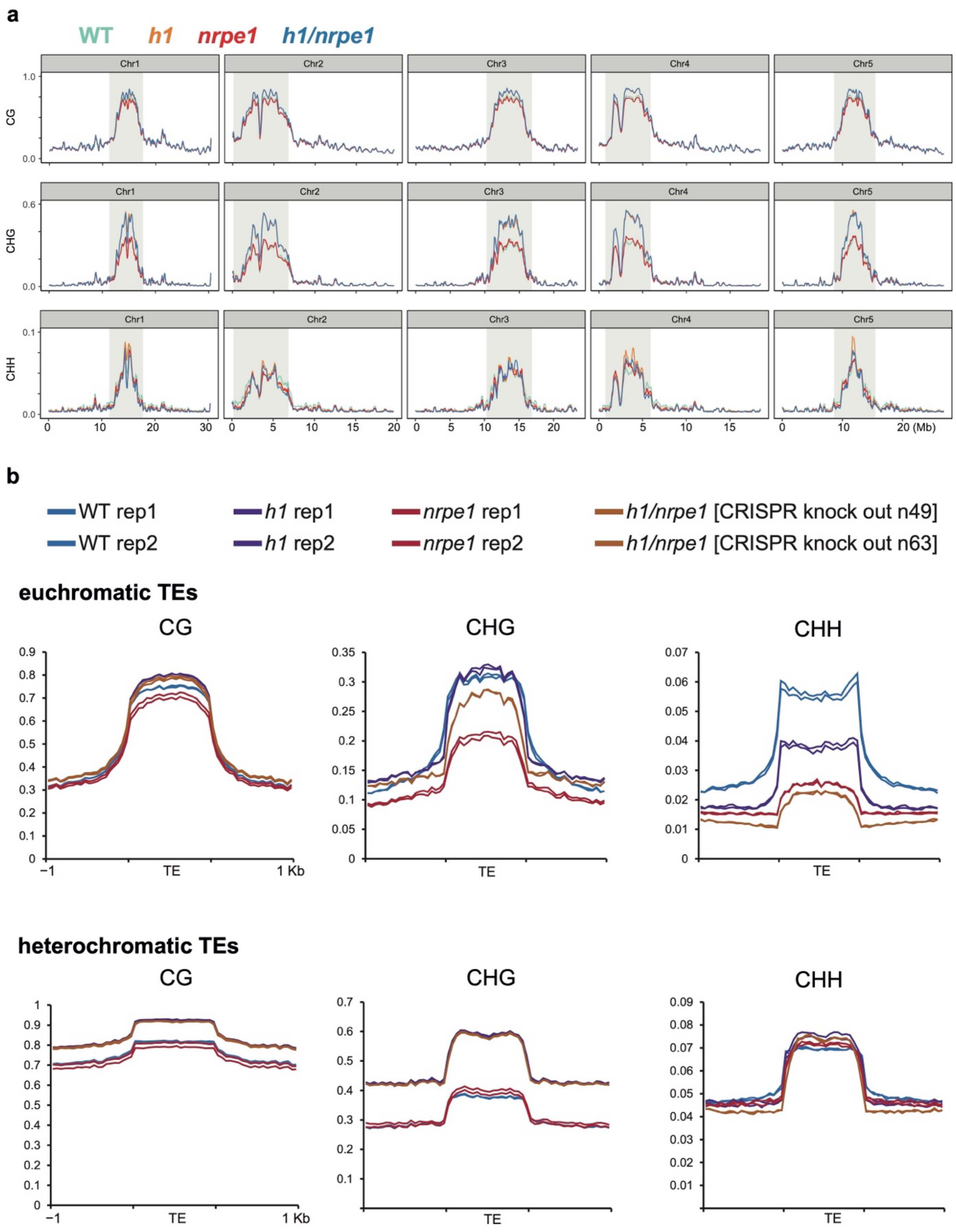


**Supplementary Figure 5:** A) Whole genome view of DNA methylation levels (average of 10kb windows) in the genotypes indicated (as in Fig 3, including all 5 chromosomes.) Y-axis indicates fraction methylation (0-1). B) Methylation metaplots over euchromatic (upper) or heterochromatic (lower) TEs (as in Figure 3, with individual biological replicates plotted separately).

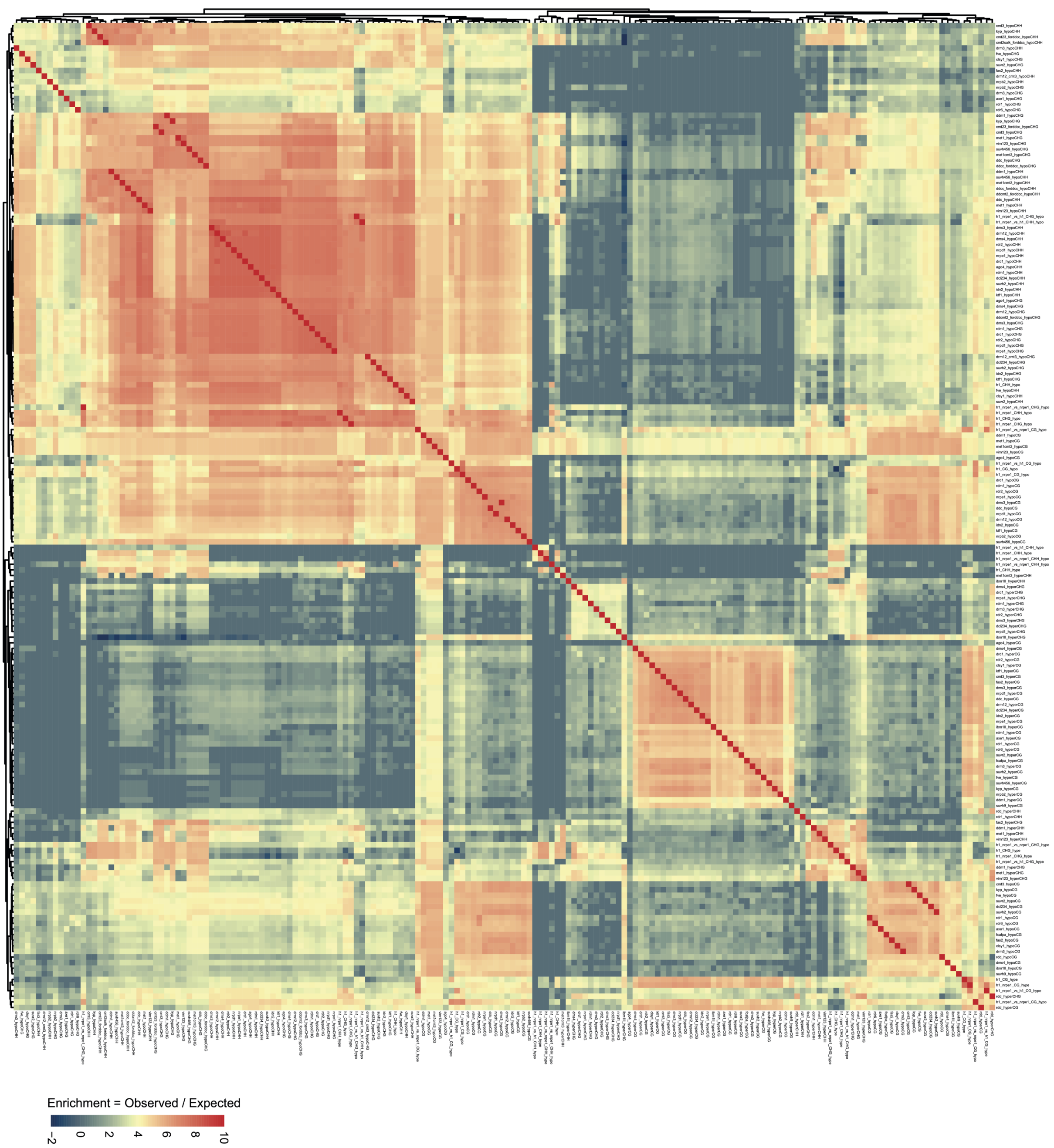


**Supplementary Figure 6:** 96 mutant genotype comparison, as shown in Figure 4, with DMR genotype comparison labels shown for visual inspection.

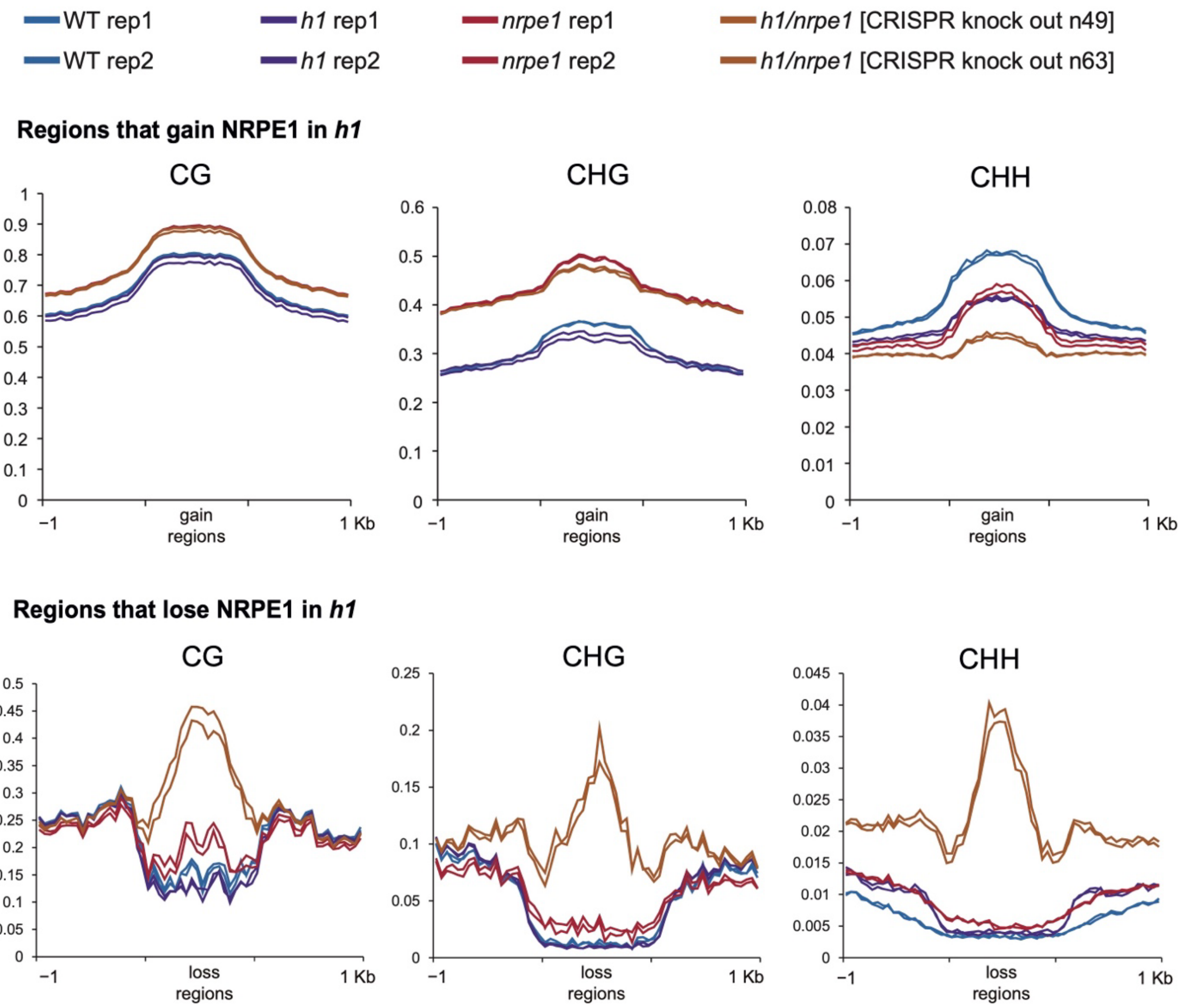


**Supplementary Figure 7**: Same data as shown in Figure 5, with individual biological replicates plotted separately.

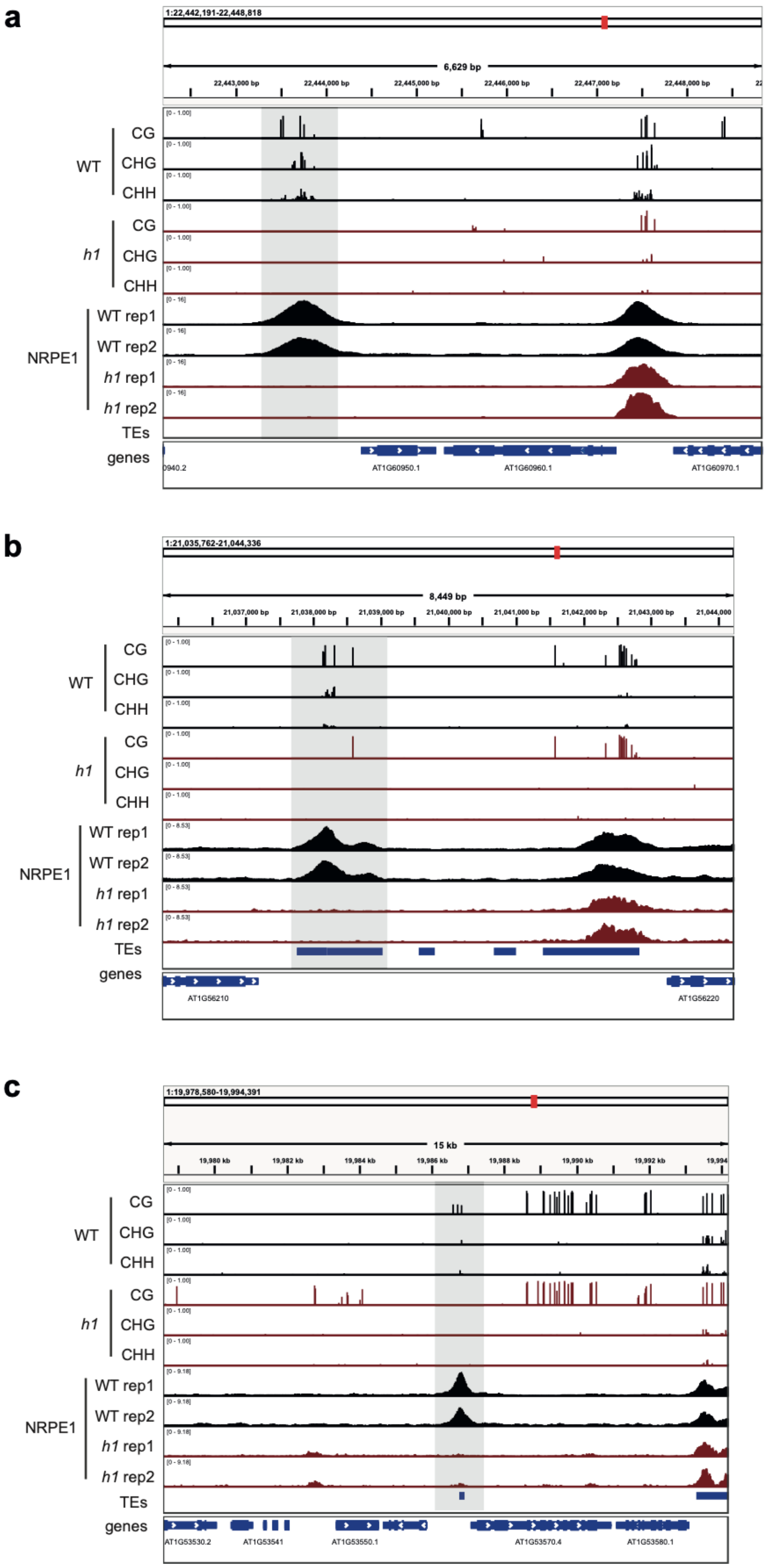


**Supplementary Figure 8:** Genome browser images of representative regions that lose both NRPE1 occupancy and DNA methylation in *h1*. Regions are highlighted in grey.
